# WSB-1 regulates DNA repair and the response to DNA damage response inhibitors in breast cancer

**DOI:** 10.1101/2025.08.27.672589

**Authors:** Effrosyni Antonopoulou, Chun Li, Amen Shamin, Scott Taylor, Flore-Anne Poujade, Michael Lind, Rajarshi Roy, Ananya Choudhury, Isabel M. Pires

**Affiliations:** Tumour Hypoxia Biology, Division of Cancer Sciences, University of Manchester, Manchester, UK; Hull York Medical School, University of Hull, UK; Translational Radiobiology, Division of Cancer Sciences, University of Manchester, Manchester, UK; Queen’s Centre for Oncology and Haematology, Castle Hill Hospital, Hull University Teaching Hospitals NHS Trust, UK; Christie Hospital NHS Foundation Trust, Manchester, UK; University of Birmingham, Birmingham, UK; CELLINK, Gothenburg, Sweden

**Author notes:** Corresponding author: Isabel M. Pires, The Oglesby Cancer Research Building, Manchester Cancer Research Centre, Wilmslow Road, Manchester, M20 4GJ, United Kingdom. These authors contributed equally to this work (Effrosyni Antonopoulou, Chun Li).

## Abstract

Tumour hypoxia is a poor prognostic factor and linked with metastatic spread and treatment resistance in most solid tumours, including breast cancer. We previously showed hypoxia-regulated E3 ligase WSB-1’s association with poor prognosis in breast cancer. Here, we evaluated the role of WSB-1 in DNA repair regulation in breast cancer, and whether these phenotypes can be exploited therapeutically. WSB-1 knockdown led to transcriptome-wide DNA repair factor upregulation, including homologous recombination (HR) factors BRCA1and RAD51. Reciprocally, WSB-1 overexpression led to downregulation of these repair factors and decreased DNA repair capacity. Patient gene expression datasets analyses also showed an inverse correlation between *WSB1* expression vs DNA repair and HR pathways. HR-deficient cancers, including BRCA1/2-deficient breast cancers, are extremely sensitive to PARP inhibitors, a phenotype described as BRCAness. We therefore hypothesised if high WSB-1 expression could be similarly exploited therapeutically. WSB-1 overexpression alone radiosensitised cells and led to increased Olaparib (PARP inhibitor) and Berzosertib (ATR inhibitor) sensitivity *in vitro*. Our study indicates that WSB-1 expression in breast cancer is associated with modulation of HR factor expression, and we propose that elevated WSB-1 expression could be considered as a potential BRCAness biomarker and promote increased sensitivity to DNA repair targeted therapy in these patients.

## Introduction

Breast cancer is the most common cancer in women worldwide [1]. Breast cancer treatment currently includes conventional therapies, such as surgery, chemotherapy, hormone therapy, and radiotherapy, as well as immunotherapy and targeted therapies [2]. Metastatic breast cancer is one of the most aggressive types of breast cancer and, unfortunately, with limited options regarding therapeutic strategies and reliable predictive biomarkers. Repression of DNA repair has been established as an inherent driver of genomic instability and tumourigenesis, as well as a potential area for therapeutic intervention. Deficiency of Homologous Recombination (HR) factors BRCA1/2 has been therapeutically exploited through its synthetic lethal interactions with PARP inhibitors [3, 4], with PARP inhibitor Olaparib recently approved for metastatic breast cancer with *BRCA1/2* germline mutations [5]. The context of HR deficiency (HRD) similar to BRCA1/2 defects, or ‘BRCAness’, can also be exploited therapeutically by targeting DDR (DNA damage response) signalling [6].

Tumour hypoxia is linked with poor prognosis in patients, including increased metastatic proneness [7]. Hypoxia is also associated with increased genomic instability linked with repressed DNA repair, including HR, and has been proposed to be therapeutically exploited in a similar manner to HR through ‘context synthetic lethality’ approaches [8].

WD repeat and SOCS box containing-1 (WSB-1) is a hypoxia inducible ECS (Elongin B/C, Cullin2/5, SOCS-box) E3 ligase [9]. WSB-1 high expression has been associated increased metastasis in several tumour types [10-15]. Our previous study has also found that WSB-1 was linked with decreased metastasis-free survival for hormone receptor-negative breast cancer patients [16]. However, its role in therapy response and DDR signalling in breast cancer remains unclear.

In this study, we aimed to evaluate the role of WSB-1 on the regulation of DNA repair in breast cancer *in vitro* models, and whether DDR inhibition in this context can be exploited therapeutically. We showed that elevated WSB-1 expression is linked with dysregulated DNA repair, and that increased WSB-1 expression sensitises to clinically relevant DDR inhibitors.

## Methods and Materials

### Cell lines, hypoxia exposure, irradiation, and drug treatment

Cell lines used in this study include MDA-MB-231 and MDA-MB-468 (triple negative mammary carcinoma), MCF7 and BT474 (Luminal A and B, respectively), and HEK-293T (immortalised embryonic kidney fibroblasts). Cells were purchased from American Type Culture Collection (ATCC) and regularly authenticated and tested negative for mycoplasma. Hypoxia exposure was performed using a H35 Hypoxystation (Don Whitley Scientific). Normoxic samples were incubated in a conventional incubator. X-ray irradiation was carried using a RS-2000 irradiator, (dose rate 1.87 Gy/min; Rad Source Technologies). HSP90 inhibitor 17-DMAG (#S1142), PARP inhibitor (PARPi) Olaparib (#S1060), and ATR inhibitor (ATRi) VE-822/Berzosertib (#S7102-SEL) (all Stratech) were prepared in DMSO and stored at −20°C.

### Plasmids and siRNA transfections

pFLAG-CMV2-WSB1 plasmid, a kind gift from Prof. Hironobu Asao, Yamagata University, Japan [17], was used for ectopic overexpression of WSB-1 using PEI (Polysciences) as previously described [18]. Transfection with non-targeting (siNT), WSB-1 (siWSB-1), and E2F-1 (siE2F1) silencing RNA (siRNA) (Supplementary Table S1) was performed using DharmaFECT#1 (Thermo Scientific) as before [16].

### qRT-PCR

RNA extraction and quantitative RT-PCR (qRT-PCR) was performed as before [19]. Analysis of relative mRNA expression was performed using QuantiFAST SYBR Green (Qiagen); primer details available in Supplementary Table S2. mRNA levels were normalised to *B2M* (β-2-microglobulin), and relative gene expression was determined using the 2^-ΔΔCT^ method [20].

### Immunoprecipitation and Immunoblotting

Whole cell lysates were prepared in UTB (Urea-Tris-Beta-mercaptoethanol) buffer as previously described [21]. Immunoprecipitation of Flag-WSB-1 and associated proteins in HEK293T cells was performed using Anti-Flag M2 Affinity Gel beads (Sigma-Aldrich) as previously described [18]. Protein levels were analysed by immunoblotting as before [21]. Antibody details are available in Supplementary Table S3. Band intensity densitometric quantification was performed using Image J (NIH) [22].

### Immunofluorescence

Immunofluorescence for DNA damage markers was performed as previously described [23], antibody details noted in Supplementary Table 4, using an Axiovert microscope (Zeiss). At least 100 foci per sample were scored using Image J.

### Flow cytometry

DNA content was analysed by flow cytometry by quantification of PI staining as before [21] and analysed using FlowJo.

### Cell proliferation/viability assays and clonogenic survival assays

The impact of drug treatments on short term cell viability was evaluated by MTS assay (CellTiter 96 AQueous One, Promega) as per manufacturer’s instructions. Clonogenic assays were performed as previously described [24], with colonies counted using the Gel Count system (Oxford Optronics). MTS viability and clonogenic survival curves were determined using a nonlinear regression curve fit of log[inhibitor] or [inhibitor] (respectively) vs response in GraphPad Prism.

### RNA-sequencing and analysis

RNA was extracted using the Aurum Total RNA mini Kit (Biorad, UK). mRNA library preparation and sequencing were performed by Novogene (Cambridge, UK), using Illumina NovaSeq PE150 platform as before [19]. Differential gene expression (DEG) analyses between siNT and siWSB-1 samples (n=3 biological replicates) were performed from read counts using the DESeq2 R package. DEGs cutoffs were adjusted p-value <0.05 and absolute fold change >log_2_(1.5). DEG volcano plots and KEGG gene ontology enrichment analyses and dotplot generation were performed as previously described [19]. Datasets will be available once study is accepted for publication.

### Mass-spectrometry analysis

Peptide analysis on FlagWSB-1 immunoprecipitated samples was performed by LC-MS/MS (liquid chromatography mass spectrometry) by the Proteomics core facility at the University of York. Peptides were matched to protein sequences using the MASCOT protein identification reference standard database. Identified proteins were ranked using peptide similarity score using log10 exponentially modified protein abundance index (emPAI) [25]. Datasets will be available once study is accepted for publication.

### Patient dataset correlation analysis

TCGA (The Cancer Genome Atlas) Breast Invasive Carcinoma (PanCancer Atlas; n = 1082) datasets were accessed through cBioportal [26, 27]. To compare *WSB1* expression against DNA repair and HR signatures, the median expression of signature genes for the corresponding molecular signatures (GSEA) was determined. Signatures log10 values were plotted against *WSB1* expression log10 values.

### Statistical analysis

All experiments included at least three independent biological replicates. Unpaired t-tests were used to compare one variable, with Holm-Šídák method used for multiple t-testing correction. Two-way ANOVA with multiple comparisons was used for multiple variables comparisons. Statistical significance was determined using Spearman’s rho rank correlation coefficients in order to examine gene expression correlations. Statistical analyses were performed using GraphPad Prism 10.6.

## Results

### WSB-1 knockdown drives extensive changes in gene expression including upregulation of DNA repair factors

To investigate the role of WSB-1 on breast cancer biology more broadly, we performed bulk RNA-sequencing analysis of transcriptomic changes upon WSB-1 knockdown (kd) in MDA-MB-231 breast cancer cells in both normoxia and hypoxia (Fig.1A-C, Supp.Fig.1A-B). We observed dramatic changes in gene expression, with pathway enrichment analysis of upregulated DEGs after kd revealing a prevalence of DNA repair relevant pathways, including homologous recombination (HR). We validated these observations in a panel of breast cancer cell lines (Fig.1D-E, Supp.Fig.1C), focusing specifically on key HR factors *BRCA1, BRCA2*, and *RAD51*, both at transcript and protein level, likely indicating WSB-1 as regulator of DNA repair generally, and HR specifically. However, the mechanism by which this occurs is not clear. We therefore sought out to identify potential factors regulated by WSB-1 that could regulate these changes.

**Figure 1.**
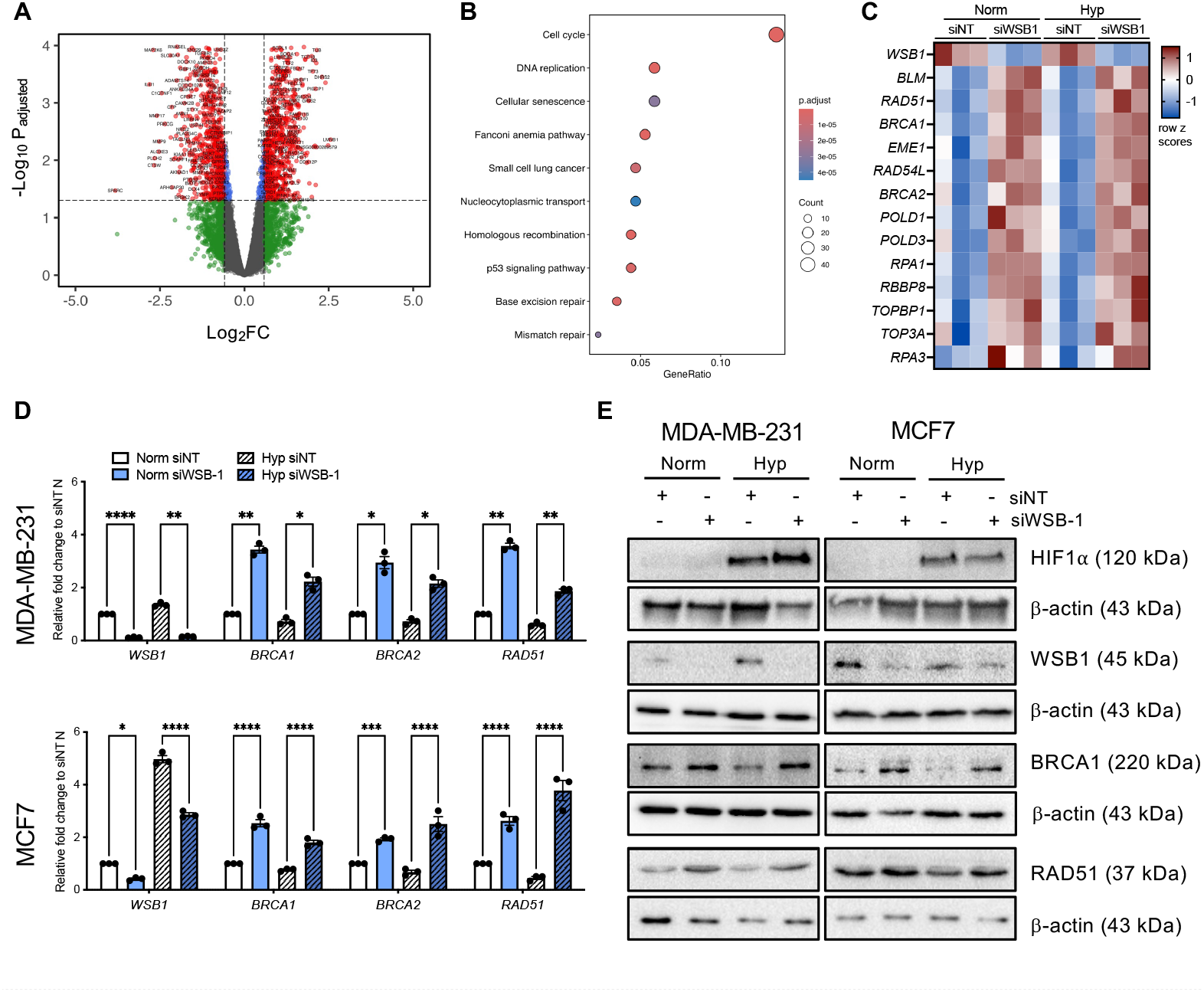
WSB-1 knockdown drives extensive changes in gene expression including upregulation of DNA repair factors. (A-C) MDA-MB-231 cells were treated with either non-targeting siRNA (siNT) or WSB-1 siRNA, (siWSB-1) exposed to hypoxic (2% O_2_) conditions for 24 hours. RNA samples were prepared and analysed by bulk RNA-sequencing (n=3). Volcano plot (A) shows significantly upregulated and downregulated genes (red) following WSB-1 knockdown. B) Enrichment plots for KEGG (for upregulated DEGs. C) Heatmap of significantly upregulated DEGs from A) from the Homologous Recombination canonical pathway (KEGG). (D-E) MCF-7 and MDA-MB-231 cells were transfected with WSB-1 siRNA (siWSB-1) or non-targeting siRNA (siNT). mRNA and protein samples were prepared after exposure to 24 h to 20% O_2_ (Norm) or 1% O_2_ (Hyp) and analysed by qPCR (D; n=3) or western blotting (E). β-actin was used as loading control and *B2M* was used as housekeeping gene. * p<0.05; ** p<0.01; *** p<0.001; **** p<0.0001. Error bars represent SDEV.

### Proteomic analysis reveals co-chaperone factors CHIP/STUB1 and HSP90 as novel interactors of WSB-1

To elucidate potential HR factor gene expression regulators downstream of WSB-1, we first performed transcription factor enrichment analysis of upregulated DEGs from our transcriptome analysis (Supp.Fig.2A). This highlighted the E2F family of transcription factors, including E2F1 and E2F4, as likely regulators of these changes. E2F1 is well established in regulating of HR factors gene expression [28, 29]. WSB-1 kd did lead to upregulation of E2F1 at transcript and protein level *in vitro* (Supp.Fig.2B). However, E2F1 kd alone did not repress *BRCA1* and *RAD51* gene expression, and E2F1 kd simultaneously with WSB-1 kd did not significantly revert the impact of WSB-1 in *BRCA1* and *RAD51* gene expression (Supp.Fig.2C). We therefore decided to analyse the WSB-1 interactome more directly to identify other potential regulators (Fig.2). For this, we first overexpressed flag-tagged WSB-1 (Flag-WSB-1), performed a pulldown of the resulting WSB-1 interacting factors, and analysed these via LC-MS/MS (Fig.2A). We were able to confirm effective WSB-1 interactor isolation by pulling down known WSB-1 ECS complex members such as Elongin B (TCEB2), Elongin C (TCEB1), and Cullin 5 (Cul5) (Fig.2A). We also isolated HSP90 isoforms and other relevant co-chaperones such as E3-ligase STUB1 (CHIP), which have been linked with DNA repair regulation [30-32]. We validated HSP90 and CHIP independently as specifically binding to WSB-1 (Fig.2B). However, when analysing the expression of these factors in whole cell lysates, their levels were not significantly altered upon WSB-1 kd (Fig.2C). We therefore predict that WSB-1 is likely binding these factors and blocking their chaperone activity but not directly targeting them for degradation. Interestingly, when HSP90 activity was blocked using HSP90 inhibitor 17-DMAG, we observed a partial reversal of the upregulation of DNA repair factor RAD51 after WSB-1 kd. However, this did not fully reverse the phenotype, indicating other factors much be involved in this process.

**Figure 2.**
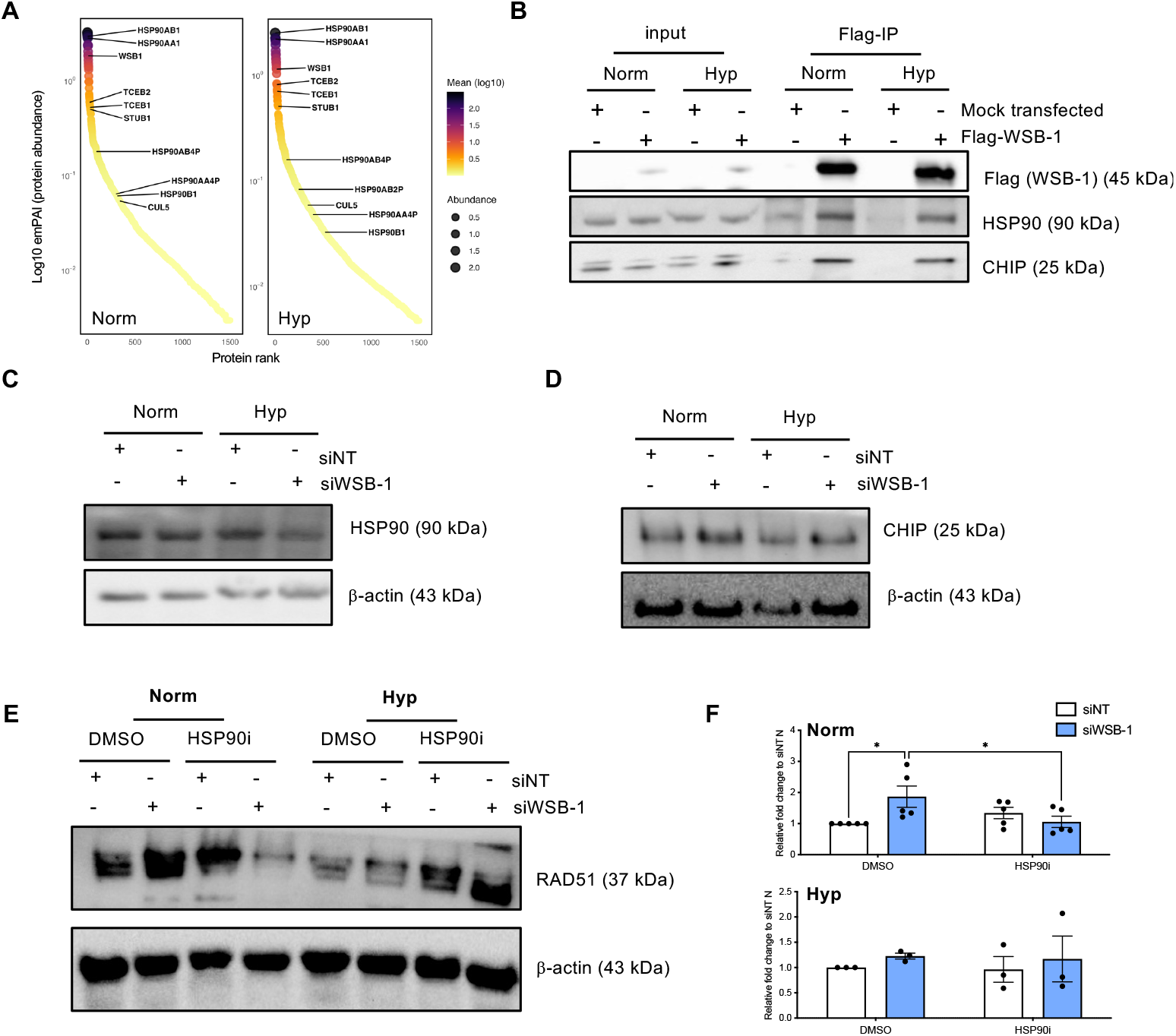
Proteomic analysis reveals co-chaperone factors CHIP/STUB1 and HSP90 as novel interactors of WSB-1. (A-B) HEK293T cells were mock transfected (mock) or transfected with a Flag-tagged WSB-1 (WSB-1) and exposed to 20% O_2_ (Norm) or 1% O_2_ (Hyp). WSB-1 binding proteins were then pulled down using Flag beads for further analysis. (A) Samples (n=3) were separated by SDS-PAGE, analysed by LC-MS/MS, and identified proteins were ranked using peptide similarity score using log10 emPAI. (B) Input and Flag-pull down (Flag-IP) samples were analysed by western blotting (n=3). (C-D) MDA-MB-231 cells were transfected with WSB-1 siRNA (siWSB-1) or non-targeting siRNA (siNT). Protein samples were prepared after exposure to 24 h to 20% O_2_ (Norm) or 1% O_2_ (Hyp) and analysed by western blotting. (E-F) MDA-MB-231 cells were transfected with WSB-1 siRNA (siWSB-1) or non-targeting siRNA (siNT) and treated with either DMSO or HSP90inhibitor 17-DMAG (HSP90i). Protein samples were prepared after exposure to 24 h to 20% O_2_ (Norm) or 1% O_2_ (Hyp) and analysed by western blotting. β-actin was used as loading control. (F) Plots represent densitometric band quantification normalised to siNT DMSO treated samples from (E). * p<0.05. Error bars represent SDEV.

### WSB-1 overexpression decreases DNA repair capacity, leading to DNA damage and cell cycle arrest

As our previous work indicated that high WSB-1 expression was associated with poor prognosis in breast cancer patients, we wanted to evaluate if WSB-1 overexpression would lead to repressed expression of genomic protective pathways such as HR. Indeed, WSB-1 ectopic overexpression led to repression of BRCA1, BRCA2, and RAD51 expression at transcript and protein level (Fig.3A-B, Supp.Fig.3). Importantly, this phenotype l was associated with decreased repair of irradiation-induced DNA damage *in vitro* (Fig.3C-D), whereas WSB-1 kd led to increased DNA repair (Supp.Fig.4A-B). Interestingly, WSB-1 overexpression led to the occurrence of DNA damage markers (*γ*H2AX and 53BPI1 foci and *γ*H2AX and KAP1 phosphorylation) in the absence of external sources (Fig.3C-E). Finally, WSB-1 overexpression led to a S-phase arrest (Fig.3F, Supp.Fig.4C). These data indicate that higher WSB-1 expression drives repression of DNA repair capacity, and the occurrence of DNA damage, triggering a DDR leading to cell cycle arrest.

**Figure 3.**
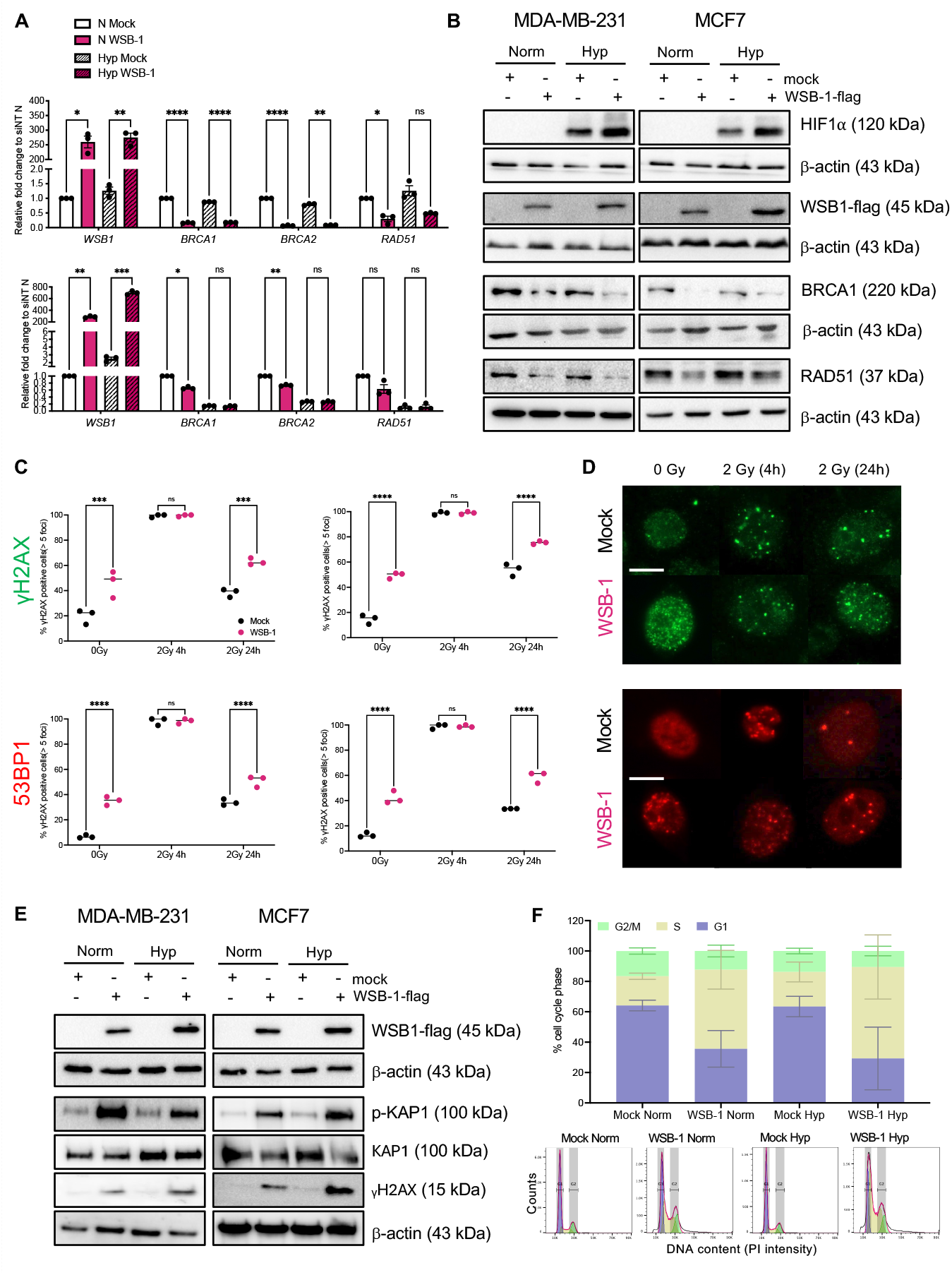
WSB-1 overexpression decreases DNA repair capacity, leading to DNA damage and cell cycle arrest. MCF-7 and MDA-MB-231 cells were mock transfected (mock) or transfected with a Flag-tagged WSB-1 (WSB-1). (A-B,E) mRNA samples and protein were prepared after exposure to 24 h to 20% O_2_ (Norm) or 1% O_2_ (Hyp) and analysed by qPCR (A; n=3) or western blotting (B, E; n=3). β-actin was used as loading control and *B2M* was used as housekeeping gene. (C-D) Presence of *γ*H2AX and 53BP1 foci were used to analyse presence of DNA damage. Dotplots (C) represent DNA repair kinetics evaluated by quantifying cells with >5 foci 4 and 24 hrs post irradiation (2 Gy) (n=3). (D) represents representative images for MDA-MB-231 cells. Scale bar = 100 *μ*m (F) MDA-MB-231 cells were mock transfected (mock) or transfected with a Flag-tagged WSB-1 (WSB-1). Cells were exposed to 24 h to 20% O_2_ (Norm) or 1% O_2_ (Hyp). Cell cycle progression was analysed by FACS (n=3). Stacked histograms represent distribution for cell cycle stages (for statistical analysis see Supplementary Figure 4); images represent examples FACs plots. * p<0.05; ** p<0.01; *** p<0.001; **** p<0.0001. Error bars represent SDEV.

### High WSB-1 expression is associated with decreased DNA repair factor expression in breast cancer patient datasets and increased sensitivity to DDR inhibitors in vitro

Our observations have been so far derived from *in vitro* studies, so we investigated if WSB-1 expression was linked with decreased DNA repair factor expression in patient samples. As predicted from our *in vitro* data, *WSB1* expression is inversely correlated to DNA repair and HR gene expression molecular signatures in breast cancer datasets (Fig.4A-B). These data, alongside our data so far, led us to hypothesise that WSB-1 overexpression could sensitise to DDR inhibitors (DDRi) as PARPi (Olaparib) and ATRi (Berzosertib), similarly to the synthetic lethal interactions between BRCA1/2 and HR deficiency and DDRi treatment. We observed that WSB-1 overexpression led to increased sensitisation to these inhibitors in both short term viability (Fig.4C) and long term clonogenic survival (Fig.4D) assays. Finally, we also observed that WSB-1 overexpression led to increased sensitisation to radiotherapy, as predicted for a DNA repair deficiency background (Fig.4E).

**Figure 4.**
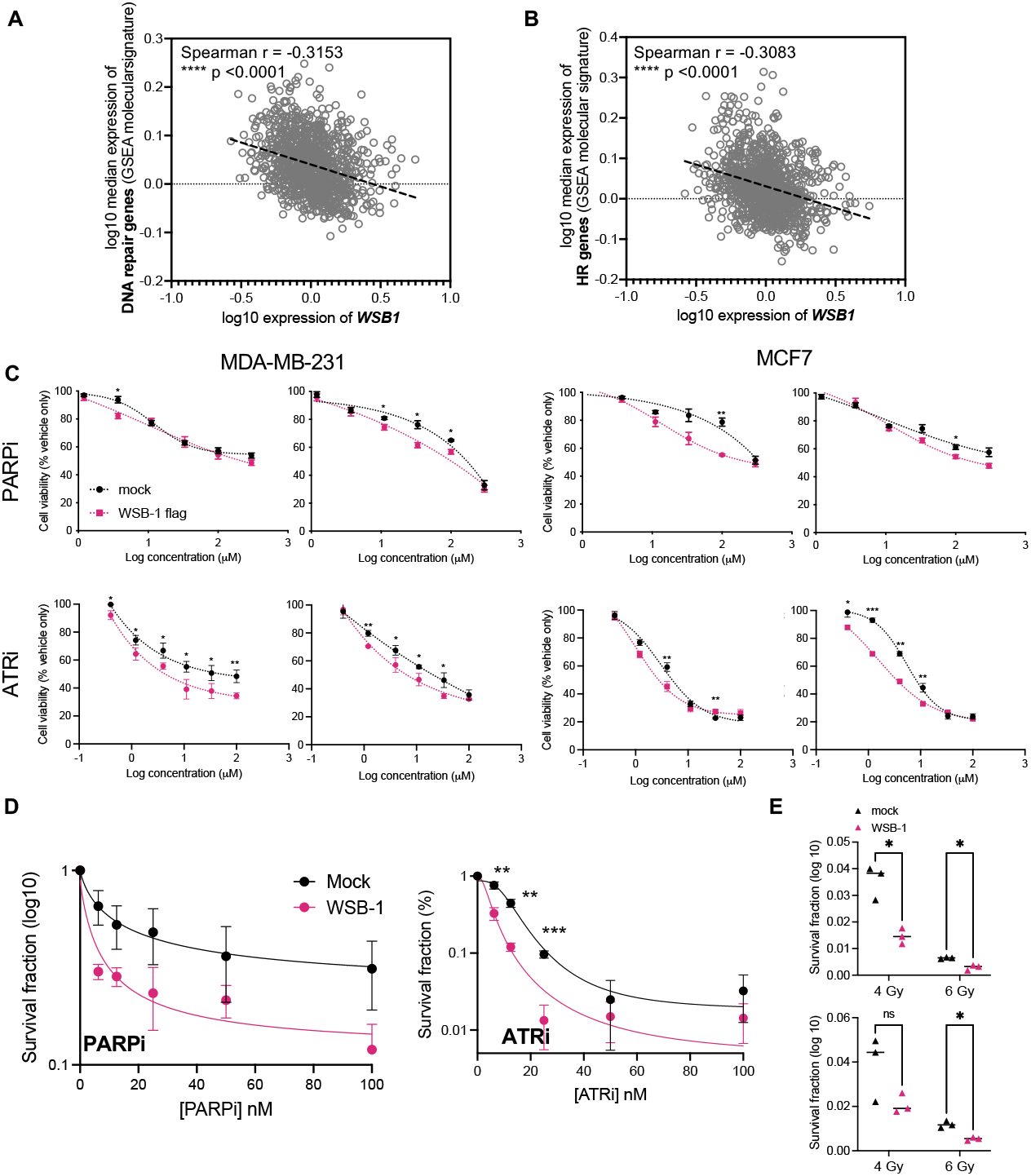
High WSB-1 expression is associated with decreased DNA repair factor expression in breast cancer patient datasets and increased sensitivity to DDR inhibitors in vitro. (A) Correlation analyses of log10 median WSB1 expression vs DNA Repair and HR (Homologous Recombination) molecular signatures (GSEA) for patient datasets in the TCGA Breast Invasive Carcinoma (PanCancer Atlas; n = 1082). (C) MCF-7 and MDA-MB-231 cells were either mock transfected (mock) or transfected with Flag-tagged WSB-1 (WSB-1). Cells were treated with a range of concentrations of PARPi (Olaparib) or ATRi (VE-822/Berzosertib) and exposed to 20% or 1% O_2_ for 72 hrs, after which MTS viability assays were performed (n=3).(D-E) Clonogenic assays for MCF-7 either mock transfected (mock) or transfected with a Flag-tagged WSB-1 (WSB-1). Single cells were treated with either vehicle (DMSO) or PARPi or ATRi as single agents (D; n=3) or RT alone or mock irradiated (E; n=3), and. Cells were left to grow into single colonies for 10 days. p<0.05; ** p<0.01; *** p<0.001. Error bars represent SDEV.

## Discussion

We previously showed that the hypoxia-regulated E3 ligase WSB-1 was linked with increased metastatic proneness in breast cancer [16]. Its role in the DNA damage response remains more elusive, beyond previous associations to aspects of the DDR through regulation of the levels of HIPK2 (linked with DNA damage-induced cell fate and cytokinesis [33]), or DDR apical kinase ATM, shown to be ubiquitylated by WSB-1 during tumour initiation and linked with oncogene-induced senescence [34]. In the present study we have demonstrated that WSB-1 modulation led to DNA repair dysregulation in breast cancer cells *in vitro*. WSB-1 kd was associated with an increase in DNA repair factor expression and reciprocally WSB-1 overexpression decreased DNA repair factors expression and reduced DNA repair capacity, alongside DNA damage occurrence. Interestingly, unlike its role in metastatic spread, which we have previously shown to be specific to hormone receptor negative breast cancer [16], WSB-1 role in DNA repair dysregulation was independent of hormone receptor status. In fact, *WSB1* expression was inversely correlated with DNA repair and HR signatures in patient samples, and increased WSB-1 expression levels led to increased sensitivity to PARP1i, ATRi, and RT.

The mechanism underpinning the impact of WSB-1 on DNA repair factors remains unclear. Initial transcriptional regulator analysis of the WSB-1 kd transcriptome upregulated DEG datasets showed E2F1 as a likely transcriptional regulator candidate downstream of WSB-1, but this was unable to be validated. Other potential transcriptional regulators downstream of WSB-1 highlighted by our study are FOXM1 and c-MYC. MYC has been linked to WSB-1 biology, with a recent study showing that WSB-1 is a direct transcriptional target of c-MYC, and that WSB-1 in turn regulates c-MYC expression via WNT/β-catenin signalling [35]. In parallel, we performed a direct analysis of the WSB-1 interactome to investigate novel WSB-1 binding partners that could impact gene expression. We identified molecular chaperones HSP90 and STUB1/CHIP as novel binding partners linked with DNA repair and gene expression regulation [30-32]. However, WSB-1 kd did not affect the overall levels of these factors, indicating any role is likely independent of WSB-1-mediate degradation of HSP90, and instead impacts its activity as a molecular chaperone. Treatment with HSP90 inhibitor 17-DMAG only partially reverse the WSB-1 kd-mediated RAD51 upregulation phenotype, indicating other yet unidentified factors are likely required for the WSB-1 mediated phenotype.

WSB-1 overexpression led to decrease in HR factors expression and repression of DNA repair capacity *in vitro*, associated with the presence of DNA damage markers even in the absence of exogenous sources of DNA damage, and an S-phase arrest. Interestingly, the type of DNA damage observed after WSB-1 overexpression indicated both more classical double strand-break γH2AX and 53BP1 foci, such as those induced by ionising radiation, but also more diffuse γH2AX staining that could indicate the presence of other types of damage, such as single strand breaks linked with replication stress [21, 23]. The presence of complex damage as well as DNA repair deficiency could possibly indicate a role of WSB-1 in driving genomic instability in breast cancer during tumour progression, to be investigated in future studies, which reinforce WSB-1’s role as an oncogene in breast cancer.

Our data indicate high WSB-1 expression could indicate HR deficiency status in tumours, as shown by the inverse correlation of *WSB1* vs DNA repair and HR gene expression signatures in patient samples, and therefore, might be amenable as a therapeutically exploitable context. *In vitro* overexpression of WSB-1 did lead to sensitisation PARP and ATR inhibition. Recently, ATRi and PARPi were shown to synergistically kill tumour cells in other HDR and DDR defective contexts in genome-wide CRISPR-Cas9 screens [36]. Therefore, the role of WSB-1 overexpression on sensitisation ATRi and PARPi combinations should also be further evaluated. PARP inhibition has been implemented as treatment strategies for a subset of BRCA1/2 and HRD deficient tumours. Whereas more work is needed to fully evaluate the mechanism underpinning WSB-1 mediated DNA repair dysregulation, high WSB-1 expression, associated with metastatic breast cancer likeliness in breast cancer, is likely to represent a proportionally larger number of patients than those with genomically-defined HDR, indicating the potential role of WSB-1 as a biomarker for response to PARP and ATR inhibitors in breast cancer, with potential to improve patient outcomes.

## Acknowledgments and funding

IMP, EA, ST, AS, and AC are funded by CRUK RadNet Manchester [C1994/A28701]. CL and FAP were supported by University of Hull PhD studentships to IMP. RNA-sequencing and proteomics interactome analysis were funded by a Breast Cancer Now (2013NovSP213) grant to IMP. The authors would like to thank Prof. Hironobu Asao (Yamagata University, Japan) for the kind gift of the pFLAG-CMV2-WSB1 plasmid and Prof Ester Hammond for helpful discussions and feedback.

## Author contributions

Project conceptualisation: IMP. Data generation and analysis: EA, CL, AS, ST, FAP. Supervision: IMP, AC, RR, ML. Writing – original draft: IMP. Writing – review & editing: all authors

## Conflicts of interest

The authors have no conflicts to declare.

## Notes

### Competing Interest Statement

The authors have declared no competing interest.

