## Supplementary material for "WSB-1 regulates DNA repair and the response to DNA damage response inhibitors in breast cancer"

**Supplementary Table S1 - List of siRNAs used in this study**

| Target gene | Catalogue number | Type and Supplier |
|---|---|---|
| Non-targeting | D-001210-02-20 | siGENOME non-targeting siRNA #2 (Horizon Discovery) |
| *WSB1* | M-013015-01-0020 | siGENOME Human WSB1 SMARTpool (Horizon Discovery) |
| *E2F1* | M-003259-01-0005 | siGENOME Human E2F1 SMARTpool (Horizon Discovery) |

**Supplementary Table S2 - List of primers used for qPCR in this study**

| Target gene symbol | Catalogue reference | Supplier |
|---|---|---|
| ***B2M*** | QT00088935 | Qiagen |
| ***BRCA1*** | QT00039305 | Qiagen |
| ***BRCA2*** | QT00008449 | Qiagen |
| ***E2F1*** | QT00016163 | Qiagen |
| ***RAD51*** | QT00031493 | Qiagen |
| ***WSB1*** | QT00064127 | Qiagen |

**Supplementary Table S3 – Immunoblotting Antibodies used in this study**

| Target | Manufacturer | Reference or Catalog No. | Dilution | Origin | Expected band size (kDa) |
|---|---|---|---|---|---|
| **β-Actin** | Santa Cruz | sc-47778 | 1:2000 | Mouse | 43 |
| **BRCA1** | Novus | NB100-598SS | 1:500 | Mouse | 220 |
| **CHIP/STUB1** | Santa Cruz | Sc-133066 | 1:1000 | Mouse | |
| **E2F1** | Cell Signalling | 3742 | 1:1000 | Rabbit | 70 |
| **Flag** | Sigma | F3165 | 1:2000 | Mouse | N/A |
| **γH2AX** | Merck Millipore | JBW301 | 1:2000 | Mouse | 15 |
| **HIF-1α** | BD Biosciences | 610958 | 1:500 | Mouse | 120 |
| **HSP90α/β** | Santa Cruz | sc-1055 | 1:1000 | Goat | 90 |
| **KAP1** | Abcam | ab10484 | 1:1000 | Rabbit | 100 |
| **Phospho-KAP1 (Ser 824)** | Cell Signalling | 4127 | 1:1000 | Rabbit | 100 |
| **RAD51** | Santa Cruz | sc-398587 | 1:1000 | Rat | 37 |
| **RAD51** | Abcam | ab63801 | 1:1000 | Rabbit | 37 |
| **WSB-1** | Cusabio | CSB-YP897589HU | 1:1000 | Rabbit | 45 |

**Supplementary Table S43 – Immunofluorescence Antibodies used in this study**

| Target | Manufacturer | Reference or Catalog No. | Dilution | Origin |
|---|---|---|---|---|
| 53BP1 | Sigma | F3165 | 1:500 | Mouse |
| γH2AX | Novus | NB100-904 | 1:1000 | Rabbit |

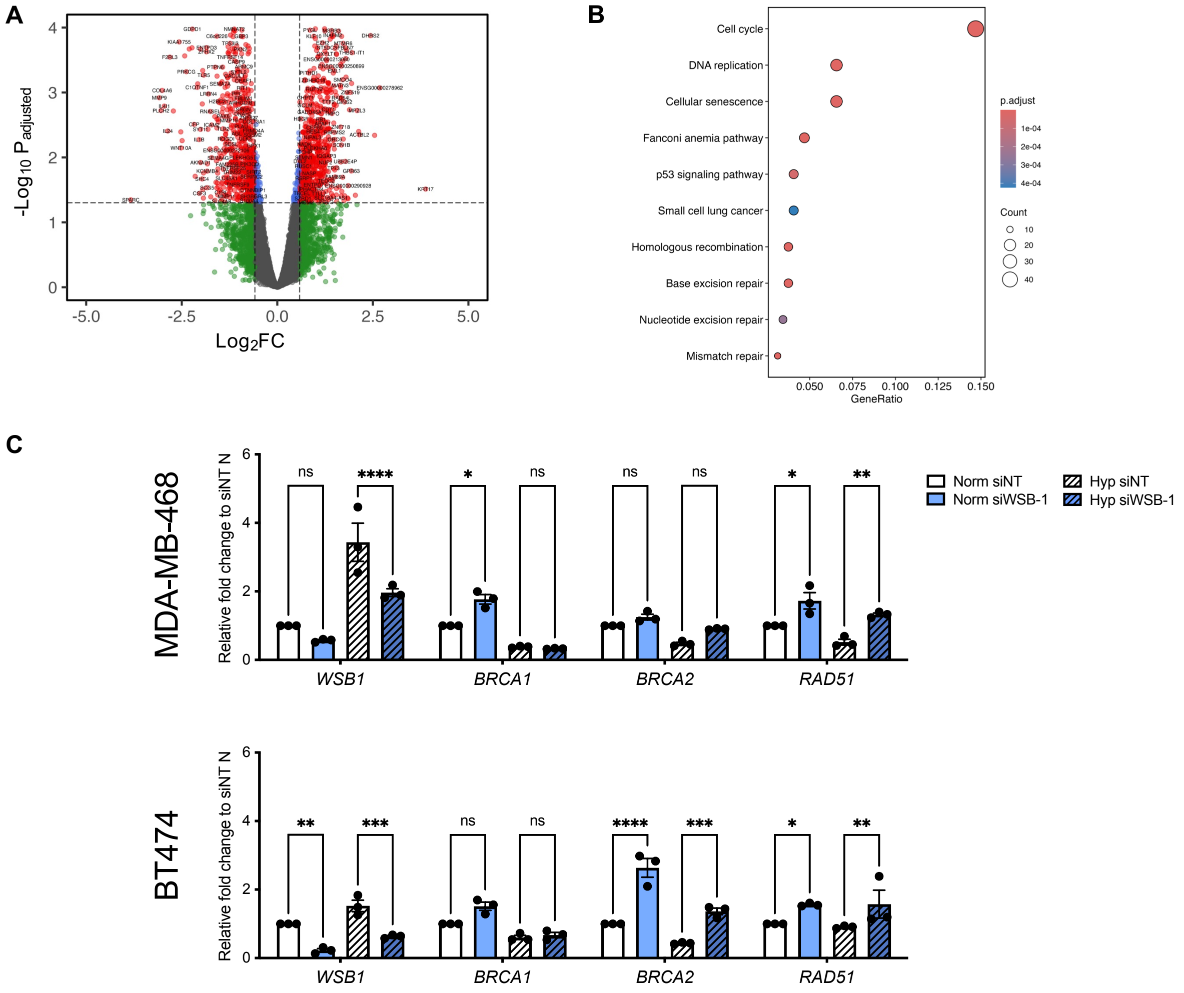


***Supplementary Figure S1 – WSB-1 knockdown drives extensive changes in gene expression and upregulates gene expression of DNA repair factors in vitro***

(A-C) MDA-MB-231 cells were treated with either non-targeting siRNA (siNT) or WSB-1 siRNA, (siWSB-1) exposed to normoxic (20% $O_2$) conditions for 24 hours. RNA samples were prepared and analysed by bulk RNA-sequencing (n=3). Volcano plot (A) shows significantly upregulated and downregulated genes (red) following WSB-1 knockdown. B) Enrichment plots for KEGG (for upregulated DEGs. (C) MDA-MB-468 and BT474 cells were transfected with WSB-1 siRNA (siWSB-1) or non-targeting siRNA (siNT). mRNA and protein samples were prepared after exposed 24 h to 20% $O_2$ (Norm) or 1% $O_2$ (Hyp) and analysed by qPCR (D; n=3). *B2M* was used as housekeeping gene. * p<0.05; ** p<0.01; *** p<0.001**** p<0.0001. Error bars represent SDEV.

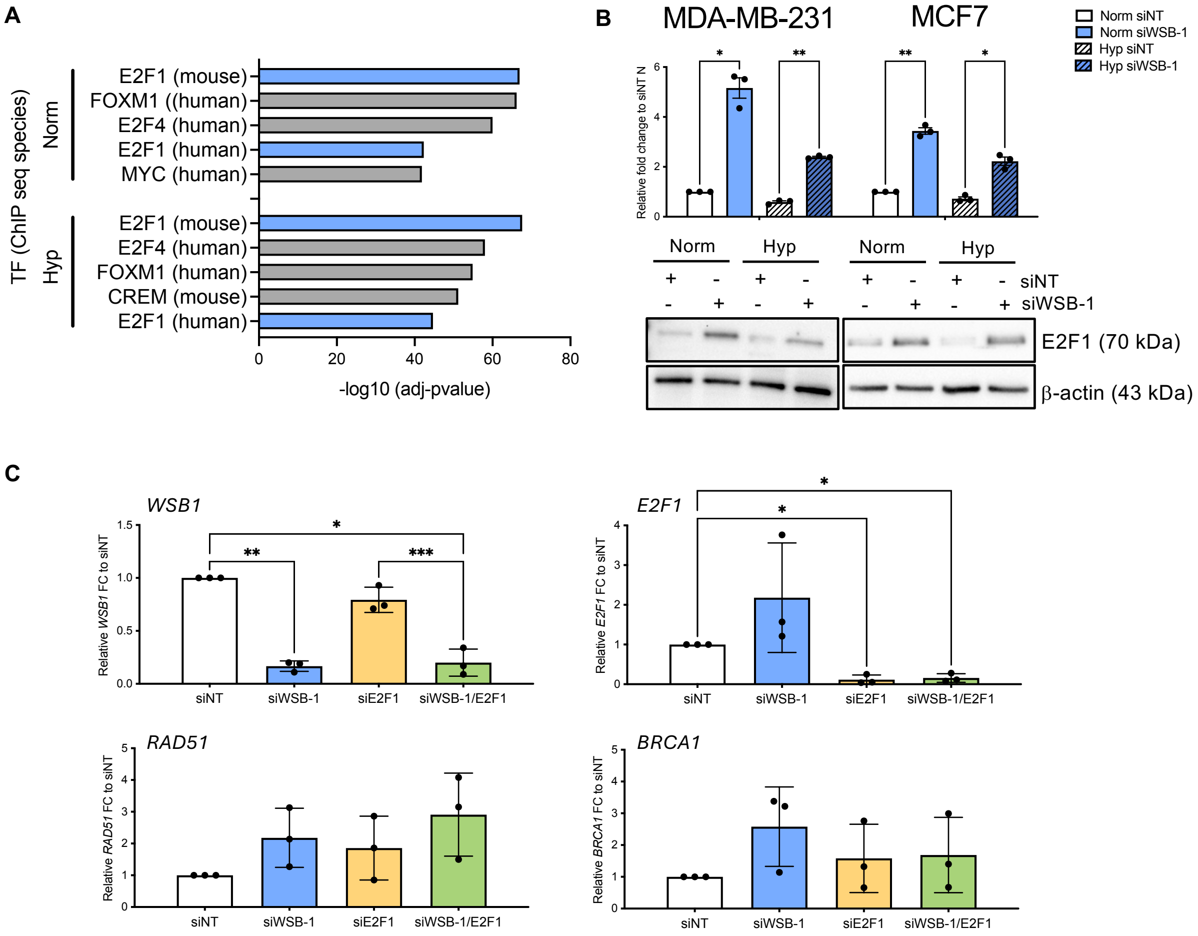


***Supplementary Figure S2 – WSB-1 knockdown impacts E2F1 expression in vitro, but does not regulate the observed changes in gene expression***

(A) Histogram represents transcription factor (TF) enrichment for the upregulated DEGs in the RNA-sequencing datasets analysed in Figure 1A-C and Supplementary Figure 1. (B) MCF-7 and MDA-MB-231 cells were transfected with WSB-1 siRNA (siWSB-1) or non-targeting siRNA (siNT). mRNA and protein samples were prepared after exposed 24 h to 20% $O_2$ (Norm) or 1% $O_2$ (Hyp) and analysed by qPCR (n=3) or western blotting (n=3). (C) MDA-MB-231 cells were transfected with non-targeting siRNA (siNT), WSB-1 siRNA (siWSB-1), E2F-1 siRNA (siE2F1), or both siWSB1 and siE2F1. mRNA samples were prepared after exposure to 24 h to 20% $O_2$ (Norm) or 1% $O_2$ (Hyp) and analysed by qPCR (n=3). *B2M* was used as housekeeping gene. * p<0.05; ** p<0.01; *** p<0.001; Error bars represent SDEV.

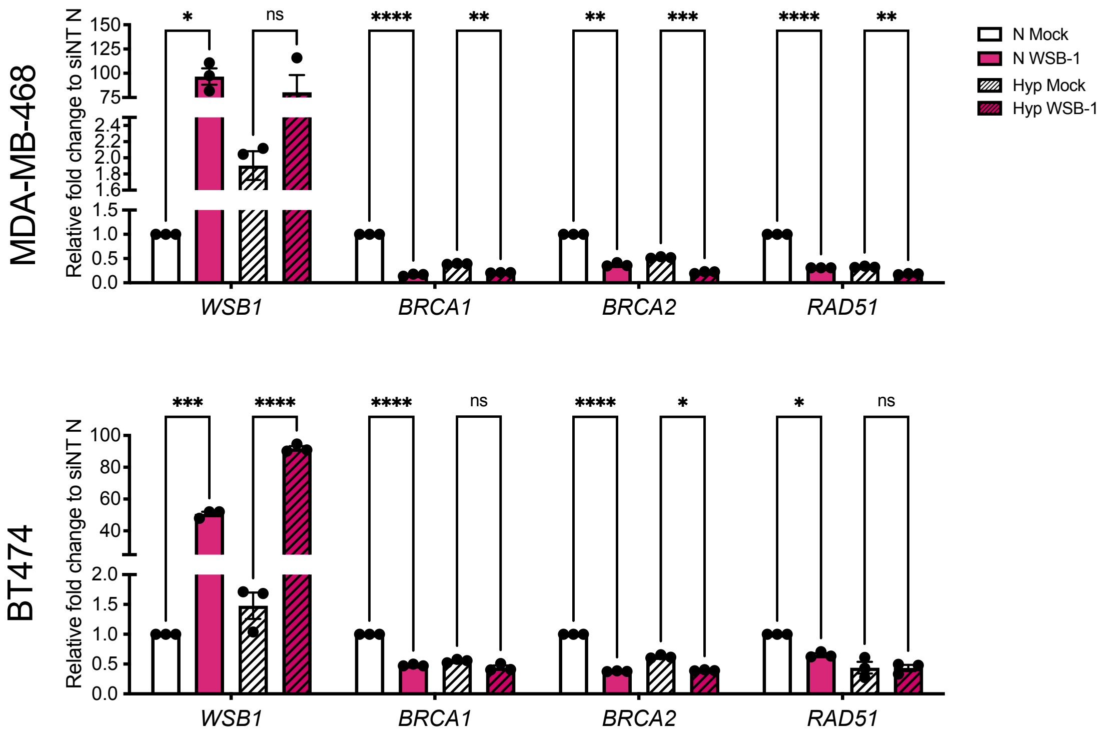


***Supplementary Figure S3 – WSB-1 overexpression drives downregulation of gene expression of DNA repair factors in vitro***

MDA-MB-468 and BT474 cells were transfected with WSB-1 siRNA (siWSB-1) or non-targeting siRNA (siNT). mRNA and protein samples were prepared after exposed 24 h to 20% $O_2$ (Norm) or 1% $O_2$ (Hyp) and analysed by qPCR (D; n=3). *B2M* was used as housekeeping gene. * p<0.05; ** p<0.01; *** p<0.001; **** p<0.0001. Error bars represent SDEV.

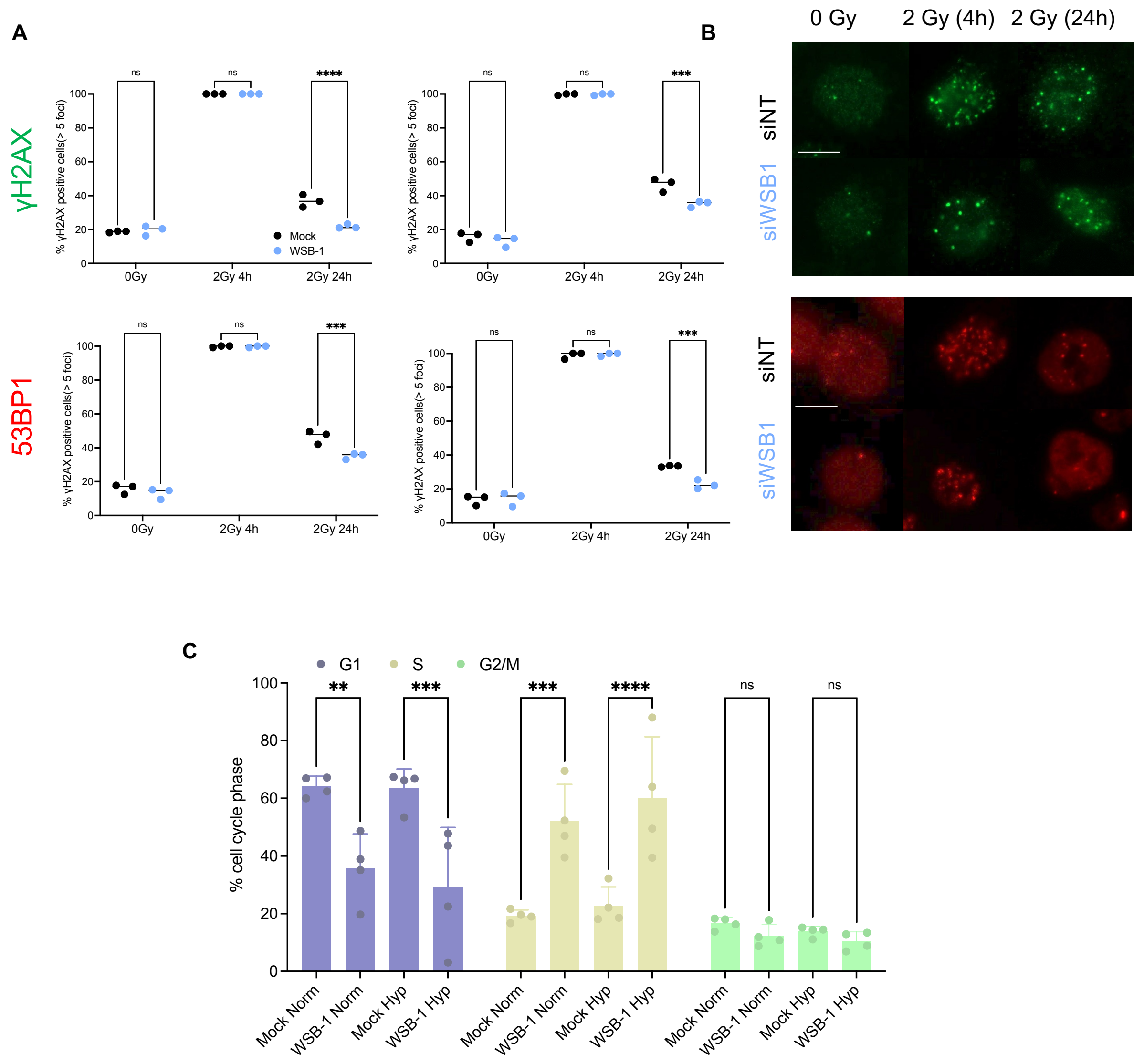


***Supplementary Figure 4. WSB-modulation impacts DNA repair capacity in vitro and WSB-1 overexpression leads to cell cycle arrest***

(A-B) MCF-7 and MDA-MB-231 cells were transfected with WSB-1 siRNA (siWSB-1) or non-targeting siRNA (siNT). Presence of γH2AX and 53BP1 foci were used to analyse presence of DNA damage. Dotplots (CA) represent DNA repair kinetics evaluated by quantifying cells with >5 foci 4 and 24 hrs post irradiation (2 Gy) (n=3). (B) represents representative images for MDA-MB-231 cells. Scale bar = 100 μm (C) Statistical significance analysis for data in Figure 3F. Histograms represent distribution for cell cycle stages. ** $p<0.01$; *** $p<0.001$****; $p<0.0001$. Error bars represent SDEV.
